# Genetic and Epigenetic Fine Mapping of Complex Trait Associated Loci in the Human Liver

**DOI:** 10.1101/432823

**Authors:** Minal Çalışkan, Elisabetta Manduchi, H. Shanker Rao, Julian A Segert, Marcia Holsbach Beltrame, Marco Trizzino, YoSon Park, Samuel W Baker, Alessandra Chesi, Matthew E Johnson, Kenyaita M Hodge, Michelle E Leonard, Baoli Loza, Dong Xin, Andrea M Berrido, Nicholas J Hand, Robert C Bauer, Andrew D Wells, Kim M Olthoff, Abraham Shaked, Daniel J Rader, Struan FA Grant, Christopher D Brown

## Abstract

Deciphering the impact of genetic variation on gene regulation is fundamental to understanding common, complex human diseases. Although histone modifications are important markers of gene regulatory regions of the genome, any specific histone modification has not been assayed in more than a few individuals in the human liver. As a result, the impacts of genetic variation that direct histone modification states in the liver are poorly understood. Here, we generate the most comprehensive genome-wide dataset of two epigenetic marks, H3K4me3 and H3K27ac, and annotate thousands of putative regulatory elements in the human liver. We integrate these findings with genome-wide gene expression data collected from the same human liver tissues and high-resolution promoter-focused chromatin interaction maps collected from human liver-derived HepG2 cells. We demonstrate widespread functional consequences of natural genetic variation on putative regulatory element activity and gene expression levels. Leveraging these extensive datasets, we fine-map a total of 77 GWAS loci that have been associated with at least one complex phenotype. Our results contribute to the repertoire of genes and regulatory mechanisms governing complex disease development and further the basic understanding of genetic and epigenetic regulation of gene expression in the human liver tissue.

## INTRODUCTION

The liver has a central role in detoxification of endogenous and exogenous toxins, synthesis of essential proteins, and regulation of carbohydrate, lipid, and drug metabolism. As such, the liver is associated with a diverse range of clinically important human traits^1^ and was recently reported as one of the most critical tissues for explaining cellular mechanisms at loci revealed by genome wide association studies (GWAS)^2^. GWAS have been effective at providing robust, but imprecise, information about genetic risk factors of complex human diseases^3^. These studies have revealed that most variation associated with complex human phenotypes do not alter protein-coding sequences, making causal variant and causal gene identification a considerable challenge^4^. Characterization of the regulatory functions of non-coding regions is the first key step towards linking non-coding regions to disease biology. Large-scale efforts such as the Encyclopedia of DNA Elements (ENCODE)^5^ and the NIH Roadmap Epigenomics^6^ consortia have made major contributions to this end. It remains an important priority to obtain similar data across many individuals, characterize the extent of between-individual variation in the activity of non-coding regulatory regions, and to identify the genetic determinants of such differential activity. Discovering genotype-dependent non-coding functional activity can help to fine map and reveal mechanisms underlying complex trait associations^7–14^. Performing such studies at genome-wide scale in large numbers of human tissues is challenging^15^ and therefore has been limited to those performed in easily accessible lymphoblastoid cell lines or blood cell types^16–22^. Here, we quantify non-coding regulatory activity in the human liver and integrate these findings with genome-wide gene expression data collected from the same human liver tissues, high resolution promoter-focused chromatin interaction maps collected from human liver-derived HepG2 cells, and GWAS summary statistics for twenty commonly studied phenotypes with variable levels of suggested causality manifesting in the liver^2^. We identify 2,625 genes and 972 non-coding regulatory elements with genotype-dependent activity in the human liver and fine-map a total of 77 GWAS loci that have been associated with at least one complex phenotype. Overall, we provide a unique resource that contributes to basic understanding of genetic and epigenetic regulation of gene expression in the human liver tissue and highlight the benefits of integrating multiple cellular traits for the identification and characterization of disease-causing genes, regulatory regions, and variants.

## RESULTS

### Annotation of non-coding regulatory regions and their interacting gene promoters

H3K4me3 and H3K27ac are epigenetic histone modifications that are enriched in functional non-coding regions of the human genome, including active gene promoters and enhancers^23–25^. We performed chromatin immunoprecipitation (ChIP)^26^ for H3K4me3 and H3K27ac modifications in human liver tissue, sequenced ChIP-ed DNA, and obtained H3K4me3 and H3K27ac ChIP-Seq data from 9 and 18 individuals, respectively (Figure 1A and Table S1, See Methods for all cohort and analysis details). Using these data, we annotated 68,600 and 131,293 genomic regions enriched for H3K4me3 and H3K27ac modifications (i.e., ChIP-Seq peaks; Figure 2A and Table S2). Similar to previous reports^23–25^, we showed that H3K4me3 is highly enriched near transcription start sites (TSS) and H3K27ac is approximately equally present within intronic, distal intergenic, and near-TSS regions (Figure 2B). We also collected genome-wide genotype and RNA-Seq data in the liver from two cohorts and, alongside with publicly available GTEx v6p liver data^27^, we studied the extent of inter-individual variation in liver gene expression levels in a total sample size of 241 (Figure 1A). Across the three cohorts in our study, we identified 23,271 expressed genes. Using a novel, high resolution, promoter focused, Capture-C based approach, termed SPATIaL-Seq (genome-Scale Promoter-focused Analysis of chromaTIn Looping) (approach was derived from^28^; see Methods for details), we identified 29,328 significant^29^ promoter-H3K4me3 and 40,839 promoter-H3K27ac peak interactions within the liver-derived HepG2 cell line (Figure 1D and Table S3). The promoters of expressed genes were significantly more likely to have DNA looping interactions with the histone peaks identified in our study relative to the promoters of genes that were not expressed (P-value <2.2×10^−16^; Figures 2C and 2D). Furthermore, the interactions between expressed gene promoters and the liver histone peaks were significantly stronger than those observed between histone peaks and genes that were not expressed (P-value < 2.2×10^−16^; Figures 2E and 2F). More than 99% of the histone peaks with evidence of looping were within 1 Mb of their interacting promoters (Figure 2G).

**Figure 1.**
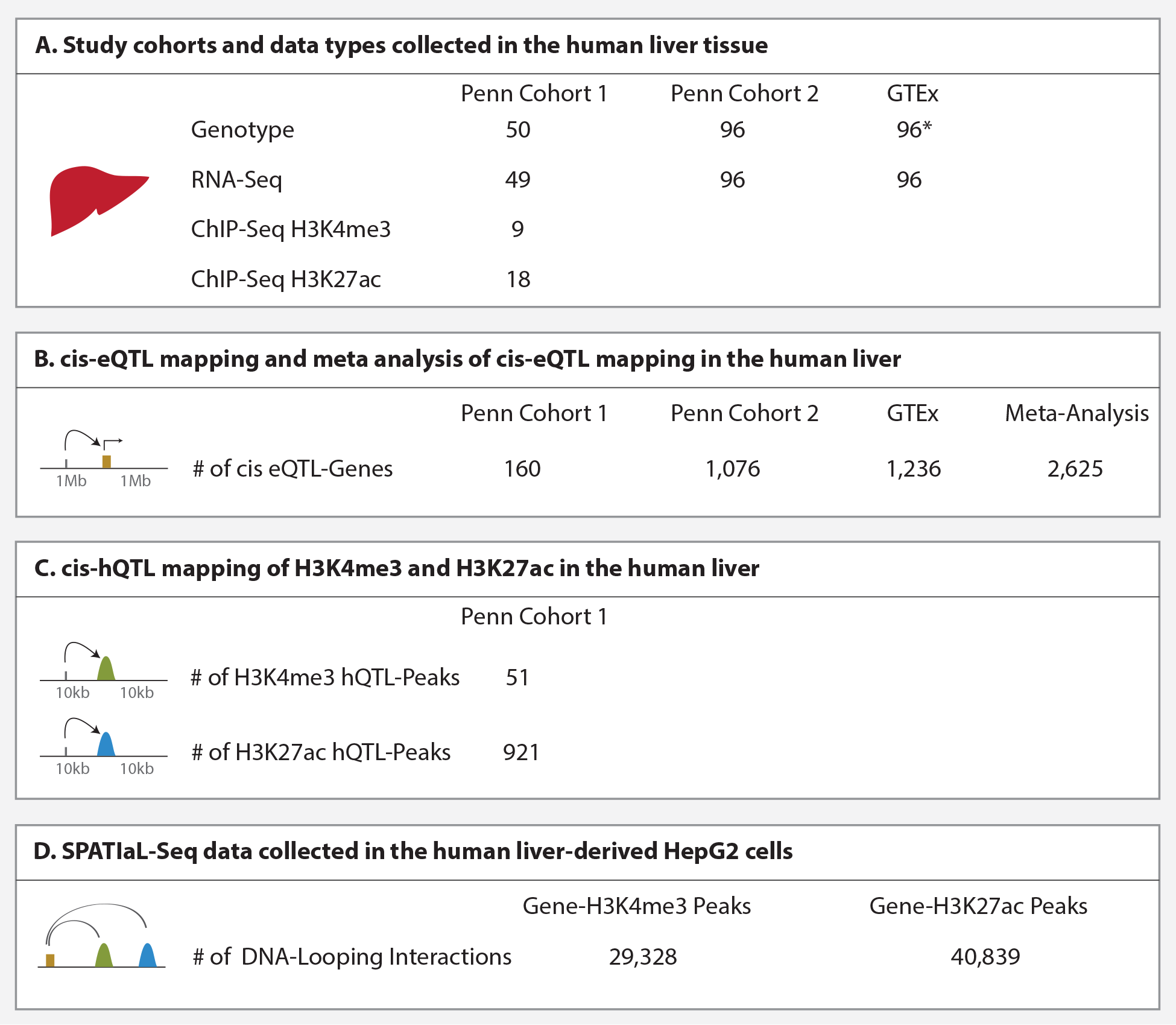
**A)** Subjects from three cohorts were included in this study. Penn cohort 1 and Penn cohort 2 samples were collected at the Penn Transplant Institute and datasets from these two cohorts have not been published previously. GTEx liver samples were collected as a part of the GTEx Analysis ReleaseV6p^27^. **B)** For eQTL mapping, associations between variant-gene pairs that are within 1 Mb of the TSS were considered as cis. cis-eQTLs were mapped within each cohort and a meta-analysis was performed across cohorts. Numbers of genes with a significant cis-eQTL at an FDR of 5% are shown. **C)** For hQTL mapping, associations between histone peaks and variants within 10 kb of the nearest end of the histone peaks were considered as cis. Numbers of histone peaks with a significant cis-hQTL at an FDR of 5% are shown. **D)** A novel, high-resolution, promoter-focused Capture-C based approach (SPATIal-Seq) was used to identify gene promoter-histone peak interactions. 29,328 gene promoter-H3K4me3 peak and 40,839 gene promoter-H3K27ac peak interactions were identified at CHiCAGO score^29^ of ≥5. *DNA from GTEx liver samples were extracted from whole blood.

**Figure 2.**
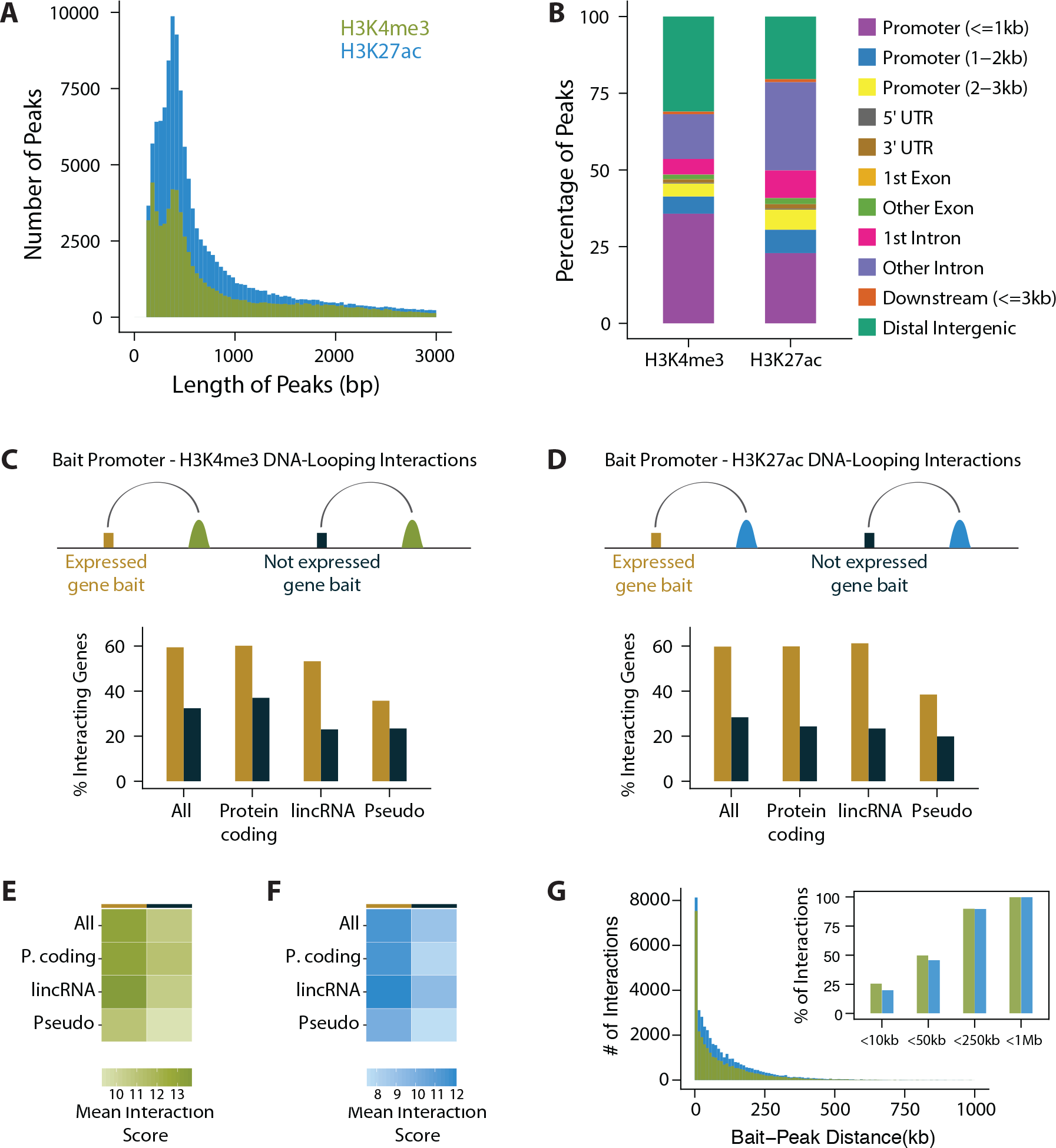
**A)** Distribution of H3K4me3 and H3K27ac peak lengths. Median peak length was 507 bp and 491 bp for H3K4me3 and H3K27ac peaks, respectively. Note that this zoomed-in plot does not display peaks with length >3,000 bp. **B)** Genomic annotations of the 68,600 H3K4me3 and 131,293 H3K27ac ChIP-Seq peaks. **C)** Percentage of baited gene promoters that form DNA-looping interactions with H3K4me3 peaks. Genes that were detected as expressed were significantly more likely to form DNA-looping interactions with H3K4me3 peaks. Pearson’s chi-squared test P-values were <2.2×10^−16^ for all, protein-coding, lincRNA gene groups and 0.00036 for pseudogenes. **D)** Percentage of baited gene promoters that form DNA-looping interactions with H3K27ac peaks. When expressed genes were compared with genes that were not detected as expressed, Pearson’s chi-squared test P-values were <2.2×10^−16^ for all, protein-coding, lincRNA gene groups and 4. 2×10^−8^ for pseudogenes. **E)** Mean CHiCAGO interaction scores between baited gene promoters and H3K4me3 peaks. Interaction scores between H3K4me3 peaks and expressed genes were significantly higher than those between H3K4me3 peaks and genes that were not detected as expressed. One-tailed Welch two sample t-test P-values were <2.2×10^−16^, 5.34×10^−6^, 0.0061, 0.084 for all, protein-coding, lincRNA, and pseudogenes, respectively. **F)** Mean CHiCAGO interaction scores between baited gene promoters and H3K27ac peaks. Interaction scores between H3K27ac peaks and expressed genes were significantly higher than those between H3K27ac peaks and genes that were not detected as expressed. One-tailed Welch two sample t-test P-values were <2.2×10^−16^, <2.2×10^−16^, 6.3×10^−5^, 8.6×10^−4^ for all, protein-coding, lincRNA, and pseudogenes, respectively. **G)** Distribution of distance between interacting bait promoters and histone peaks. >99% of all interacting bait-H3K27ac peak pairs were within less than 1 Mb apart.

### Identification of shared and liver-specific cis-eQTLs

We mapped cis-expression quantitative trait loci (cis-eQTL; here defined as associations between the expression level of a gene and a variant within 1Mb of the gene TSS) within each cohort by linear regression^30^ and performed a meta-analysis^31^ across three cohorts. We identified 2,625 genes with cis-eQTLs at 5% FDR; we hereafter refer to such genes as eQTL-Genes (Figure 1B and Table S4). For each cis-eQTL identified, we estimated the posterior probability^32^ that the eQTL effect is present in 43 non-liver GTEx v6p tissues^27^ (Figure 3A and Table S4). We classified cis-eQTLs that have a posterior probability of greater than 0.9 for being an eQTL in at least 38 non-liver tissues as “shared eQTLs” and those with a posterior probability of greater than 0.9 in fewer than five non-liver tissues as “liver-specific eQTLs” (the last and first quartiles of the distribution in Figure 3A, respectively). We integrated these cis-eQTL findings with H3K4me3 and H3K27ac peaks that we identified in the human liver as well as those that were identified in multiple cell lines by the ENCODE consortium^5^ (Table S5). Overall, cis-eQTLs were significantly more likely to overlap H3K4me3 and H3K27ac histone peaks relative to randomly selected SNPs matched for key properties including linkage disequilibrium (LD), minor allele frequency (MAF), gene density, distance to nearest gene (Figure 3B and Table S6). Moreover, shared eQTLs overlapped H3K4me3 promoter marks more often than liver-specific eQTLs (Figure 3B and Table S6). Conversely, while shared eQTLs showed similar levels of overlap with H3K27ac enhancer marks across different tissues, liver-specific eQTLs were significantly more likely to overlap H3K27ac marks that we identified in human liver tissue as well as those that were identified in liver-derived HepG2 cell lines, consistent with the previous reports of significantly higher cell-type specificity of enhancers relative to promoters^33^ (Figure 3B and Table S6). Despite implementing multiple approaches, we did not identify any significant trans-eQTLs in the human liver (See Methods for details).

**Figure 3.**
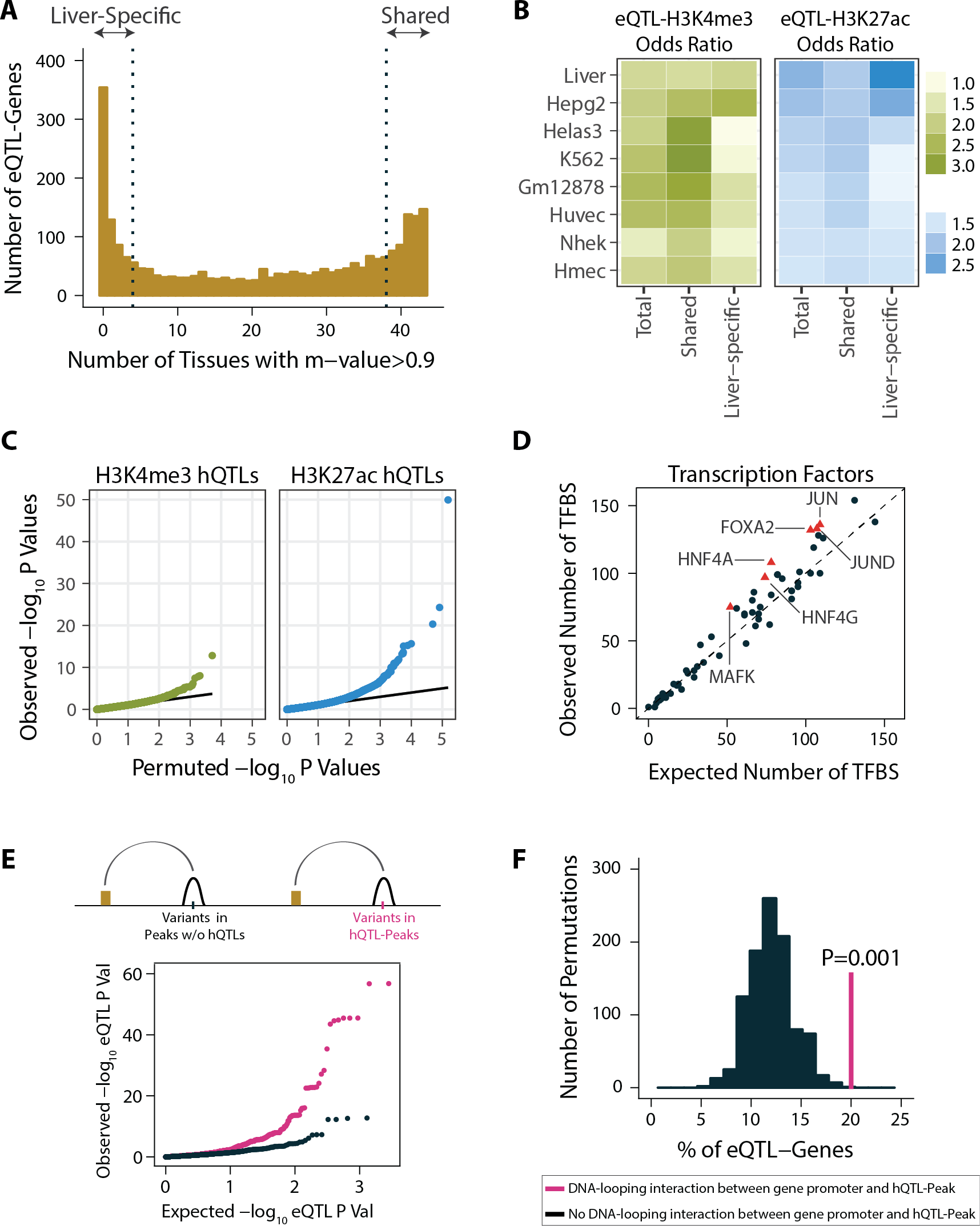
**A)** Distribution of the number of non-liver GTEx tissues with an association posterior probability (m-value) of >0.9 for lead cis-eQTL-Gene pairs. cis-eQTL-Gene pairs in the first quartile of the distribution were considered as liver-specific, those in the last quartile were considered as shared eQTLs. **B)** Overlap between cis-eQTL sets (total, shared, and liver-specific) and H3K4me3 and H3K27ac peaks. Odds ratios are relative to randomly chosen matching (with respect to LD, MAF, gene density, distance to nearest gene) SNP sets. H3K4me3 and H3K27ac data from non-liver tissues were obtained from ENCODE database, links to the ENCODE data files are included in Table S5. The odds ratios are only plotted for cell lines with both H3K4me3 and H3K27ac data. Results from other cell types are included in Figure S13, P-values and odds ratios are included in Table S6. **C)** QQ-plots of the cis-hQTL association P-values of H3K4me3 (Panel 1) and H3K27ac (Panel 2). Solid lines represent the expected distribution of P-values based on permuted data. **D)** Observed and expected numbers of transcription factor binding sites (TFBS) in hQTL-Peaks. Expected numbers represent the mean TFBS overlap of 1,000 set of randomly chosen matching numbers of autosomal liver histone peaks. Transcription factors that were significantly enriched (One-tailed Fisher’s exact test P-value <0.05) in hQTL-Peaks are shown in red triangles. **E)** hQTL-Peaks were assigned to their interacting gene(s) using chromatin capture data. For each SNP within a hQTL-Peak, its eQTL-Pvalue on the interacting gene(s) is plotted in magenta color. Matching numbers of peaks were drawn from the set of expressed and baited genes that do not interact with hQTL-Peaks. eQTL P-values of the SNPs within the background set of histone peaks on their interacting gene(s) are shown in black color. One-sided Wilcoxon rank sum test P-value comparing two P-value distributions was 3.75×10^−9^. **F)** Percentage of eQTL-Genes among genes that interact with hQTL-Peaks is shown in the magenta vertical line. Distribution of expected percentage based on 1,000 sets of randomly chosen matching numbers of genes that do not interact with hQTL-Peaks are shown in black. P-value of 0.001 is based on the permutation test^93^.

### Identification of cis-hQTLs in the human liver

To identify genetic determinants of H3K4me3 and H3K27ac modifications in the liver, we applied a method that uses both total and allele-specific signals in sequencing data to enable quantitative trait loci (QTL) mapping with relatively small sample sizes^20^. We identified cis-QTLs for 51 H3K4me3 and 921 H3K27ac peaks at 5% FDR (Figures 1C, 3C and Table S7). We refer to such variants as histone QTLs (hQTLs) and the peaks that they regulate as hQTL-Peaks throughout the manuscript. We intersected the hQTL-Peaks with transcription factor binding sites (TFBS) that were obtained in HepG2 cells by the ENCODE consortium (Table S5)^5^. We found that liver hQTL-Peaks are significantly enriched for binding sites of hepatocyte nuclear factors (HNF4A, HNF4G, FOXA2) as well as transcription factors (TF) involved in hepatocellular remodeling (JUN and JUND) when compared with randomly selected matching numbers of liver histone peaks from our data (P-value <0.05; Figure 3D). Further, using our chromatin capture data, we found that variants within hQTL-Peaks were more likely to be significantly associated with the expression of genes with which they are physically in contact with relative to the variants within histone peaks that do not have hQTLs (P-value = 3.75×10^−9^, Figure 3E). Overall, genes that physically interact with an hQTL-Peak were almost twice as likely to have cis-eQTLs relative to randomly selected matching numbers of expressed and baited genes that do not interact with hQTL-Peaks (P-value = 0.001; Figure 3F). These results suggest that genotype-dependent putative functional elements identified here play causal roles in the regulation of gene expression levels and this, at least in part, is mediated via DNA looping interactions.

### Co-regulated histone modification states and gene expression levels

Integrating eQTL associations with cis-regulatory element annotations has proven useful for the precise identification of causal regulatory variants and the specific cis-regulatory elements they perturb. Our results highlight the value of analyzing cell-type-matched gene expression and cis-regulatory element datasets (Figure 3B). These analyses, however, are limited as there are often multiple candidate regulatory regions within each eQTL locus and hence it has remained difficult to systematically link regulatory elements to their respective target genes. To address this, we first identified putatively co-regulated hQTL-Peaks and eQTL-Genes. Because of the limited sample size of our ChIP-Seq data, we identified co-regulated Peak-Gene pairs based on LD between lead QTL-SNPs (i.e., r^2^>0.8 between the lead hQTL and lead eQTL; Figure 4A). We found 116 Gene-Peak pairs that are likely regulated by the same causal variant (Table S8). These 116 Gene-Peak pairs corresponded to 104 unique eQTL-Genes and 95 unique hQTL-Peaks. hQTL-Peaks were often not assigned to their nearest gene; in 71% of the co-regulated Gene-Peak pairs, there was at least one other gene that is closer to the hQTL-Peak than the eQTL-Gene with which it is co-regulated (Figures 4B and 4C). Figure 4D displays an example of a putatively genetically co-regulated Gene-Peak pair, supporting the presence of a shared causal effect underlying the activity of an enhancer located in the second intron of the *SPIRE1* gene (H3K27ac-84963; chr18: 12,551,731-12,553,678) as well as *SPIRE1* gene expression level. The full set of putatively genetically co-regulated Gene-Peak pairs are included in Table S8.

**Figure 4.**
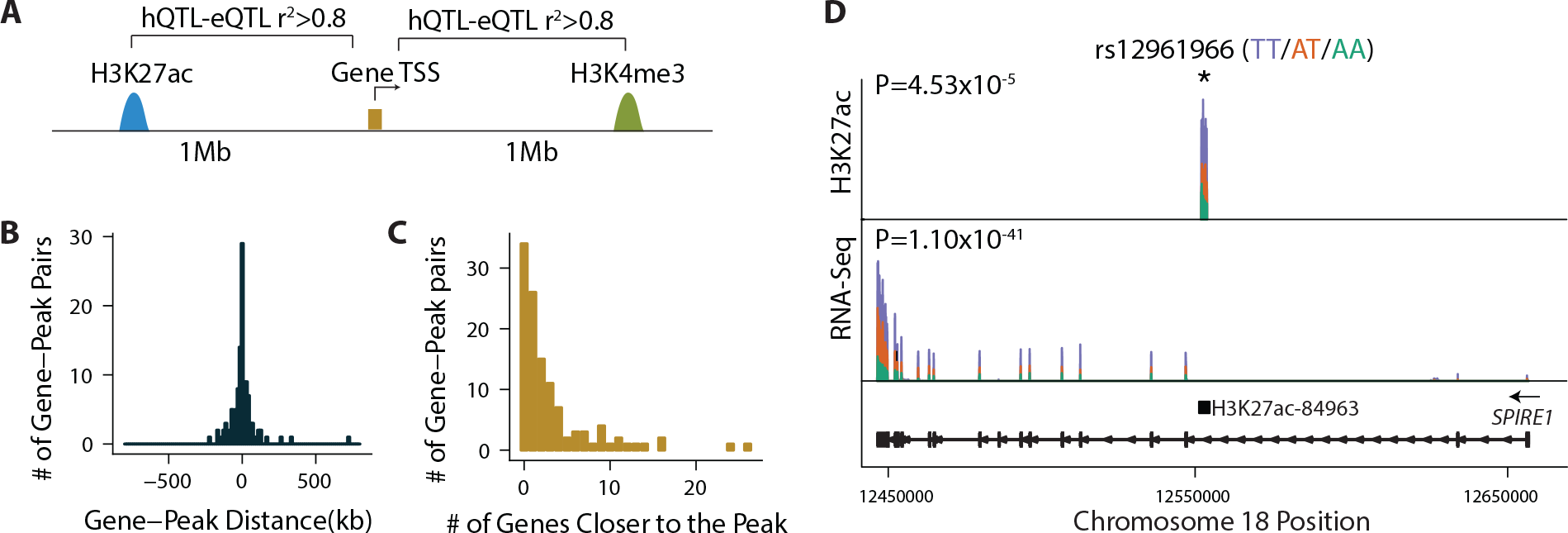
**A)** For each gene with a significant meta cis-eQTL, H3K4me3 and H3K27ac peaks with significant hQTLs that are located within 1 Mb of its transcription start site (TSS) were selected as candidate co-regulated peaks. Gene-Peak pairs with r^2^>0.8 between the lead hQTL and lead eQTL were considered as putatively co-regulated. **B)** Distance between co-regulated Gene-Peak pairs. **C)** Distribution of number of genes that are closer to the hQTL-Peak than its putatively co-regulated eQTL-Gene. **D)** Example of a putatively co-regulated Gene-Peak pair. SNP rs12961966 was significantly associated with chromatin modification state of an enhancer (H3K27ac-84963; chr18: 12,551,731^−12^,553,678) residing in the second intron of the *SPIRE1* gene and *SPIRE1* gene expression level. Sushi plots^94^ show the mean normalized read counts of each genotype group. Sample sizes of each genotype group were TT:4, AT:8, AA:5 for ChIP-Seq data and TT:55, AT:94, AA:88 for RNA-Seq data. *SPIRE1* gene model shown below the sushi plots was generated using ggbio Bioconductor package^95^ and *SPIRE1* transcript ENST00000409402.

### Identification of candidate causal genes and gene regulatory regions in GWAS loci

Next, we asked whether leveraging our hQTL, eQTL, and chromatin capture findings could help fine-map GWAS loci. Throughout this manuscript, we defined “fine-mapping” as evidence of refinement in putatively causal-variant, causal-regulatory region, and causal-gene identification in any individual GWAS locus. We obtained GWAS summary statistics of twenty phenotypes that are commonly studied (based on the number of PubMed IDs in the NHGRI-EBI GWAS Catalog) and that have variable levels of suggested causality manifesting in the liver^2^. These phenotypes included coronary artery disease^34^, HDL cholesterol^35^, LDL cholesterol^35^, total cholesterol^35^, triglycerides^35^, diastolic blood pressure^36^, systolic blood pressure^36^, mean arterial pressure^36^, rheumatoid arthritis^37^, type 2 diabetes^38^, multiple sclerosis^39^, asthma^39^, psoriasis^39^, Parkinson’s disease^39^, Alzheimer’s disease^40^, schizophrenia^41^, Crohn’s disease^42^, ulcerative colitis^42^, inflammatory bowel disease^42^ and age-related macular degeneration^43^. Links to the GWAS summary statistics used are included in Table S9. We used a P-value threshold of < 1 x 10^−6^, selected a lead variant to represent each 2 Mb region (1 Mb upstream and 1 Mb downstream of the lead variant), and identified 1,614 loci previously associated with these phenotypes. Genetically regulated gene expression levels and histone modification states in GWAS loci can reveal the mechanisms underlying observed associations between genetic variants and disease phenotypes^7–14^. We therefore applied a Bayesian colocalization approach^44^ to identify eQTL-Genes co-regulated with the disease phenotypes (Figure 5A), and used an LD threshold (r^2^>0.8) between lead GWAS SNPs and lead hQTL-SNPs to identify putatively causal cis-regulatory elements in GWAS loci (Figure 5B). In colocalization analysis, we first assessed whether there was sufficient power to test for colocalization (PP3+PP4>0.8), and for the colocalization pairs that pass the power threshold, we defined the significant colocalization threshold as PP4/(PP3+PP4)>0.9^16; 44^.

**Figure 5.**
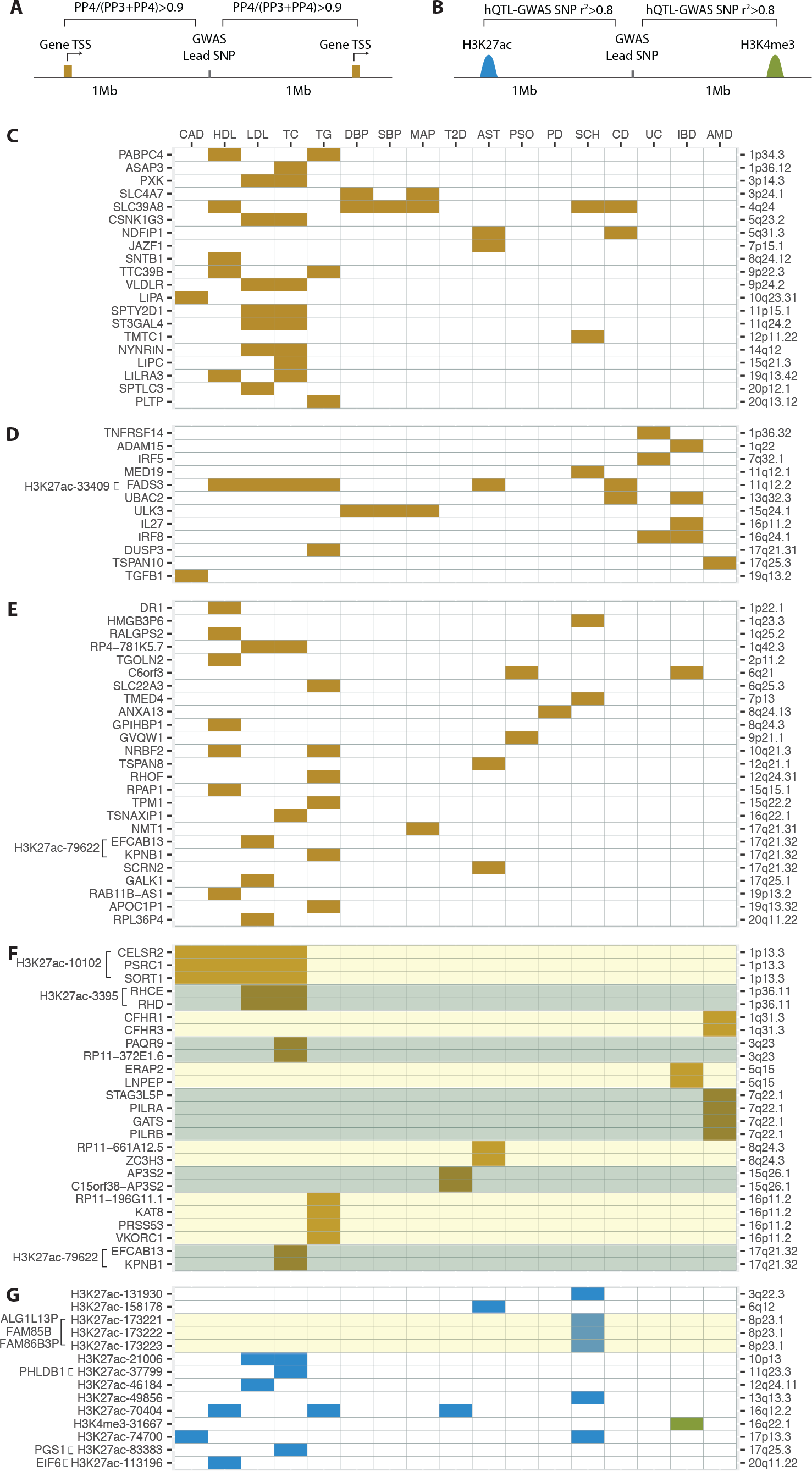
**A)** Each GWAS locus was defined as the 2 Mb region around the lead GWAS variant. A Bayesian colocalization approach was performed between the GWAS phenotype and each gene with a significant meta cis-eQTL whose TSS resides within the GWAS locus. Among colocalization pairs that pass the colocalization power threshold (PP3+PP4>0.8), the ones that met PP4/(PP3+PP4)>0.9 threshold were considered as significantly colocalized. **B)** GWAS-Peak pairs with r^2^>0.8 between the lead hQTL and lead GWAS SNP were considered as putatively co-regulated. Only hQTL-Peaks that reside within the GWAS loci were considered as co-regulation pairs. **C)** GWAS loci with only one significantly colocalized gene and the gene identified has been reported as the likely causal gene by overwhelming majority of the literature. **D)** GWAS loci with only one significantly colocalized gene and the gene identified has been reported among several candidate genes that were suggested to be causal in the literature. **E)** GWAS loci with only one significantly colocalized gene and the gene identified has not been previously implicated in the corresponding GWAS phenotype. **F)** GWAS loci with more than one significantly colocalized gene. Genes within the same GWAS locus are shown in alternating shades of yellow and blue. When identified, putatively causal histone peaks are included next to the colocalized gene names of each locus. **G)** GWAS loci where putative causal regulatory regions were identified in the absence of colocalized liver eQTL-Genes. Seven out of these 12 regions could be assigned to eQTL-Genes in non-liver GTEx tissue types and those assigned genes are included next to the histone peak IDs in the figure. Phenotype abbreviations are as follows: coronary artery disease (CAD), HDL cholesterol (HDL), LDL cholesterol (LDL), total cholesterol (TC), triglycerides (TG), diastolic blood pressure (DBP), systolic blood pressure (SBP), mean arterial pressure (MAP), Type 2 diabetes (T2D), asthma (AST), psoriasis (PSO), Parkinson’s disease (PD), schizophrenia (SCH), Crohn’s disease (CD), ulcerative colitis (UC), inflammatory bowel disease (IBD), age-related macular degeneration (AMD)

We found a total of 121 GWAS-Gene and 33 GWAS-Peak pairs with evidence of shared genetic causality (Table S10). For 85 GWAS-Gene pairs, our dataset contains evidence supporting identification of the causal gene, as there was only one gene that significantly colocalized with the GWAS phenotype (Figure 5C). We identified several genes and gene regulatory regions that underlie associations with more than one GWAS phenotype. For instance, 85 GWAS-Gene pairs with only one colocalized gene implicated 57 unique genes (Figures 5C-5E). 20 of these genes were previously reported as the likely causal gene at the locus^35; 36; 41; 45–52^ (Figure 5C). For 12 loci, our findings help refine causal gene identification from among several candidates that were suggested to be causal in the literature^36; 46; 49; 52–58^ (Figure 5D) and in 23 loci we identified novel putatively causal genes (i.e., genes that were not previously implicated with the GWAS phenotypes in our study) (Figure 5E). We were not able to identify any causal genes for rheumatoid arthritis, multiple sclerosis, and Alzheimer’s disease using the data collected in the human liver.

At the 17q21.32 locus, we identified a genetically regulated enhancer (H3K27ac-79622; chr17:45,733,609-45,733,977) that likely underlies GWAS associations with LDL cholesterol, triglyceride, and total cholesterol levels. We found *EFCAB13* and *KPNB1* as the likely causal genes driving associations with LDL and triglyceride levels, respectively (Figure 5E). We were not able to distinguish the effects of these two genes with regards to total cholesterol associations (Figure 5F). The colocalization probabilities of these two genes were close to the significance threshold for all three phenotypes, suggesting that there is insufficient signal to discriminate the three genes using colocalization analysis. The enhancer identified in this locus, however, was only forming DNA-looping interactions with the promoter of the *KPNB1* gene, and the direction of effect on enhancer activity was only consistent with *KPNB1* expression, suggesting that *KPNB1* is the likely causal gene in this locus (Figure 6A). Similary, at the chromosome 1p13.3 locus, we reassuringly identified the previously reported causal enhancer (H3K27ac-10102; chr1:109,816,977-109,818,871)^11^ as the regulatory region responsible for the GWAS associations with coronary artery disease, HDL cholesterol, LDL cholesterol, and total cholesterol levels (Figure 5F). Using our chromatin capture interaction data, we further demonstrated that this enhancer only interacts with the promoter of the *SORT1* gene in the genome, prioritizing *SORT1* as the causal gene in this locus, consistent with the previous reports^11^ (Figure 6B).

**Figure 6.**
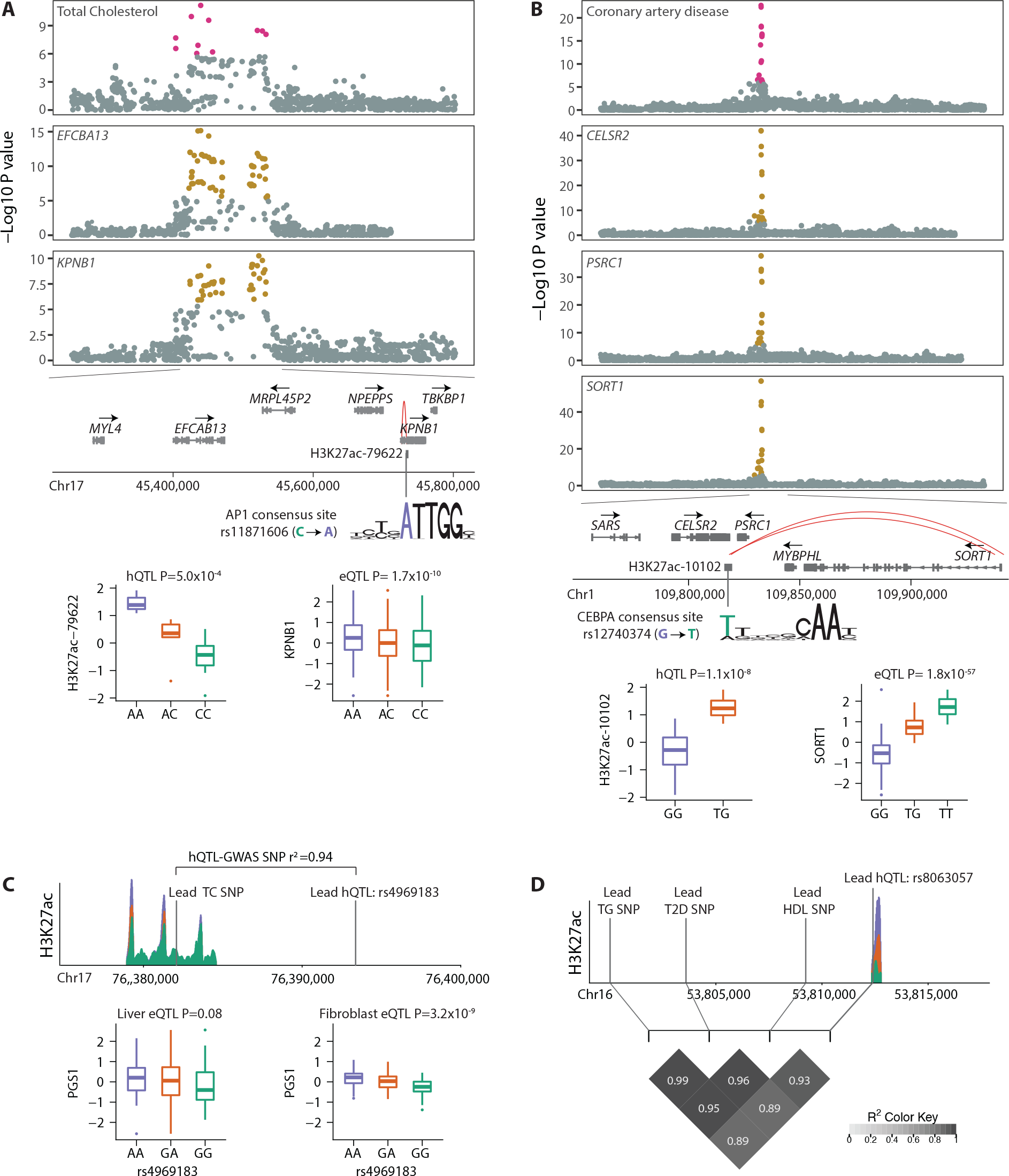
**A)** Significant colocalization signals at the 17q21.32 locus are displayed using Manhattan plots. Colocalization posterior probability of total cholesterol GWAS associations with *EFCAB13* gene expression was 0.999 and with *KPNB1* was 0.934. Schematic representation of the genes in the zoomed-in locus of chr17: 45,250,000^−45^,800,000 and the putatively causal H3K27ac-79622 peak (chr17:45,733,609^−45^,733,977). H3K27ac-79622 peak only showed significant DNA looping interaction with the promoter of the *KPNB1* gene in the genome (CHiCAGO score: 7.26). SNP rs11871606 is the lead hQTL of this peak and significantly increases the odds of AP1 binding (P=4.89×10^−4^). Box plots of normalized H3K27ac-79622 ChIP-Seq and *KPNB1* RNA-Seq read counts are stratified by genotype at the rs11871606. Sample sizes of each genotype group were AA:3, AC:5, CC:10 for ChIP-Seq data and AA:49, AC:111, CC:81 for RNA-Seq data. **B)** Significant colocalization signals at the chromosome 1p13.3 locus. Colocalization posterior probabilities of coronary artery disease associations with *CELSR2*, *PSRC1*, and *SORT1* gene expression levels were all 0.999. Schematic representation of the genes in the zoomed-in locus of chr1: 109,750,000^−109^,940,573 and the putatively causal H3K27ac-10102 peak (chr1:109,816,977^−109^,818,871). H3K27ac-10102 peak only formed DNA looping interaction with the promoter of the *SORT1* gene in the genome (CHiCAGO scores: 6.63 and 7.8). SNP rs12740374 located within this peak and increases the odds of CEBPA binding (P=0). Box plots of normalized H3K27ac-10102 ChIP-Seq and *SORT1* RNA-Seq read counts are stratified by genotype at the rs12740374. Sample sizes of each genotype group were GG:14, TG:4 for ChIP-Seq data and GG:143, TG:83, TT:14 for RNA-Seq data. **C)** Example GWAS locus with putatively causal regulatory region (H3K27ac-83383; chr17:76,378,946^−76^,384,578) identified in the absence of colocalized eQTL-Gene in the human liver. Sushi plot showing the mean normalized H3K27ac-83383 ChIP-Seq read counts of each genotype group at rs4969183. Lead hQTL and lead total cholesterol GWAS SNP are shown in vertical lines on the sushi plot. H3K27ac-83383 was assigned to a lipid metabolism gene, *PGS1*, in the transformed fibroblasts (r^2^>0.8 between lead hQTLs and lead eQTL). Boxplots show *PGS1* gene expression levels stratified by the genotype at the lead hQTL. Sample sizes of each genotype group were AA:65, GA:116, GG:56 in the human liver tissue and AA:73, GA:126, GG:64 in transformed fibroblasts. **D)** 16q12.2 GWAS locus where putatively causal regulatory region identified in the first intron of the *FTO* gene (H3K27ac-70404; chr16:53,812,377^−53^,812,817). Sushi plot show the mean normalized H3K27ac-70404 ChIP-Seq read counts of each genotype group at rs8063057. This peak could not be assigned to any gene in the human liver tissue data from our study as well as 43 non-liver tissue types of the GTEx consortium. LD heatmap show the r2 between lead hQTL SNP as well as the lead GWAS SNPs of T2 diabetes, HDL, and triglyceride levels.

At GWAS loci where putatively causal regulatory regions were identified in the absence of colocalized eQTL-Genes (Figure 5G), we examined whether these regulatory regions could be assigned to target genes in non-liver tissue types of the GTEx consortium using the approach described in Figure 4A (i.e., r^2^>0.8 between the lead hQTL and lead eQTL). In four out of twelve such loci, we were able to find the putative target genes in non-liver tissue types (Figure 5G). Among these was the chromosome 17q25.3 locus where we found a genotype-dependent enhancer (H3K27ac-83383; chr17:76,378,946-76,384,578) with evidence of underlying GWAS association signal with the total cholesterol levels. We were not able to assign this peak to any gene and found no significant colocalization signal in this locus with the total cholesterol levels in the human liver (Figure 6C). However, using the cis-eQTL data from transformed fibroblasts^27^, we assigned this peak to a lipid metabolism gene, *PGS1* (Figure 6C), suggesting that *PGS1* is the likely causal gene in this locus. This result also suggests that the downstream effects of genotype-dependent chromatin modification states might be more pronounced in certain tissues depending on the power of eQTL mapping or the presence of condition specific transcription factors, as was reported previously^16^. Lastly, while we were not able to identify a target causal gene at the chromosome 16q12.2 locus, we found a genetically regulated putative enhancer in the first intron of the *FTO* gene (H3K27ac-70404; chr16:53,812,377-53,812,817) with evidence of underlying GWAS associations in this locus with type 2 diabetes, HDL, and triglyceride levels (Figure 6D). To our knowledge, this is the first report of a genotype-dependent regulatory element identified in the human liver that overlaps the GWAS interval at this locus, which has received considerable study^59^.

### Characteristics of fine-mapped GWAS loci

For each GWAS locus with at least one candidate causal gene or regulatory region identified, we estimated the probability that an individual variant among all tested variants is causal given the pattern of QTL association at that locus^60; 61^. In ten loci, we identified fewer than five candidate causal variants accounting for 95% of the total posterior probability of being causal for that locus (Figure 7A).

**Figure 7.**
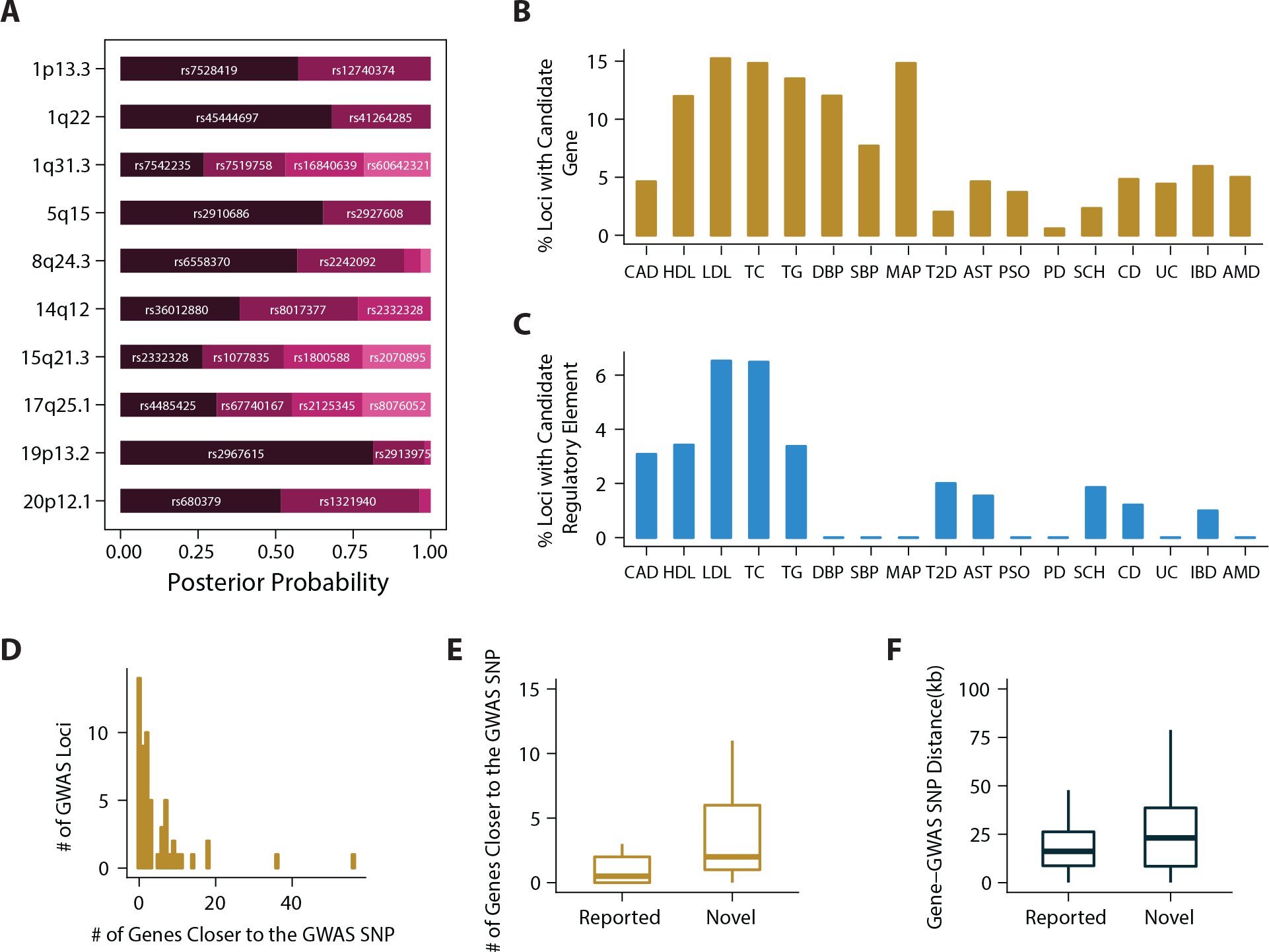
**A)** 10 loci with fewer than five candidate causal variants and the candidate causal variants with their corresponding posterior probabilities are shown. **B)** Percentages of GWAS loci with at least one significantly colocalized gene are shown for each of the complex trait phenotype. **C)** Percentages of GWAS loci with at least one candidate regulatory element are shown for each of the complex trait phenotype. **D)** Number of genes that are closer to the GWAS lead SNP than the causal gene identified. GWAS loci with only one colocalized gene are included in this figure. **E)** Number of genes that are closer to the GWAS lead SNP are shown for genes that were previously reported to be causal (i.e. genes shown in Figure 5C) and for the novel disease causing genes identified in our study (i.e. genes shown in Figure 5E). **F)** Distance between GWAS lead SNP and the TSS of the causal gene identified. “Reported” corresponds to genes that were previously reported to be causal (i.e. genes shown in Figure 5C) and “Novel” corresponds to the novel disease causing genes identified in our study (i.e. genes shown in Figure 5E).

Overall, using the genome-wide data collected in the human liver, we were able to fine-map at least one GWAS locus for 17 out of 20 phenotypes we studied (Figure 5). 31% of the candidate causal genes identified had liver-specific cis-eQTLs, 19% had shared cis-eQTLs, and the remaining 50% had cis-eQTLs that were neither classified as liver-specific nor as shared. Supporting the validity of our approach, our ability to identify causal genes and gene regulatory regions in GWAS loci depended on the physiological relevance of the studied phenotype to the human liver. Phenotypes with known, apparent causality derived from the liver had larger proportion of GWAS loci fine-mapped in our study (Figures 7B and 7C).

A median of two genes were located closer to the lead GWAS SNP than the causal gene identified in this study (Figure 7D). When we compared the novel causal genes identified in our study (Figure 5E) to genes that were previously suggested to be causal in the literature (Figure 5C), we noted a significant difference both in terms of the distance and the number of genes between the lead GWAS SNPs and the causal genes identified (Figures 7E and 7F). This discrepancy emphasizes once again that genes reported as causal in the literature are biased towards those that are closer to the GWAS lead SNPs and that unbiased genome-wide approaches are required to identify true causal genes in GWAS loci.

We also note that mutations in four of the complex phenotype-causing genes (*LIPA*, *LIPC*, *GPIHBP1*, *IRF8*) have been implicated in related Mendelian diseases^62–65^ and 52% of the causal genes that have murine models were reported to display similar phenotype in the model organism as well ^66,878787^(Table S11). Lastly, while *LIPA*, *PLTP*, and *SLC39A8* have been previously suggested to affect their associated phenotypes through protein altering mutations^67–69^, our findings suggest that genotype-dependent changes in their gene expression levels also contribute to the complex trait pathogenesis.

## DISCUSSION

In 2001, the first published draft of the human genome confirmed that the vast majority of its sequence, approximately 97 percent of the 3.2 billion bases, has no protein coding function^70^. Following this discovery, the next phase of research focused on understanding and functionally annotating non-coding regions within the human genome. These studies generated reference epigenomic maps for multiple cell lines and tissue types and demonstrated that epigenetic marks on histone proteins are important predictors of gene-regulatory activity^71^. Perhaps more interestingly, such gene regulatory regions were subsequently shown to harbor the majority of the complex disease associated variants^4^, making studies of gene regulation an important area of investigation at the interface of basic and disease biology.

In this study, we generated the most comprehensive genome-wide dataset yet of two epigenetic marks, H3K4me3 and H3K27ac, in the human liver. Using DNA-looping interactions, we identified at least one target interacting regulatory element for 65.4% of the genes that were baited and detected as expressed. We demonstrated widespread functional consequences of natural genetic variation on regulatory element activity and gene expression levels. Further, we showed that a single genetic variant could co-regulate both histone modification states and gene expression levels and this co-regulation is at least partly mediated via DNA looping interactions. We expect that this expansive resource containing functional annotation of non-coding elements and DNA-looping interactions between gene promoters and putative functional gene regulatory elements will greatly facilitate future analyses and stimulate new areas of investigation.

Our results also hold significant relevance for medical genomics. Using genetic co-regulation approaches, we fine-mapped a total of 77 GWAS loci associated with at least one complex phenotype. For 20 loci, the causal gene we identified had been previously reported as causal by the majority of the literature. For 12 loci, our findings helped refine the causal gene among several reported candidate causal genes and for 23 loci, we identified novel causal genes. For a total of 16 loci, we identified a candidate causal gene regulatory region and for 10 loci, we pinpointed the causality to fewer than five potential variants with greater than 95% certainty.

While our efforts constitute the largest GWAS fine-mapping effort performed in the human liver, we were able to identify causal genes in less than 20% of the GWAS loci even for the most directly liver related complex phenotypes (i.e. blood lipid levels). This result indicates a need for similar comprehensive studies of the transcriptome and epigenome in a wider range of tissue types and stimulation conditions as well as studies focusing on other complex disease-causing mechanisms.

Overall, our findings expand the repertoire of genes and regulatory mechanisms governing complex disease development and contribute to basic understanding of genetic and epigenetic regulation of gene expression in the human liver tissue. Furthermore, by more precisely highlighting genes and regulatory elements with relevance to disease or critical intermediate phenotypes, we believe this study will improve research into the development of therapeutic or preventative measures to mitigate the effects of complex disease. Finally, our approaches to integrate genetic variation and multiple molecular phenotypes across individuals are likely to be applicable to other tissues and traits.

## MATERIAL AND METHODS

### Study Subjects

#### Penn Cohort 1

Samples in this cohort were prospectively collected between August 2014 and February 2015 at the Penn Transplant Institute. The cohort was comprised of 50 liver donors (49 deceased, 1 living). Sex and age of the donors were reported as 19 females, 31 males aging between 6 and 77 years old.

#### Penn Cohort 2

Between January 2012 and August 2014, ~25 mg of liver needle biopsy samples were collected from deceased donors prior to transplantation surgery at the Penn Transplant Institute. All samples were stored in RNAlater. For this study, 96 samples were chosen based on cold ischemic time (138 – 320 minutes) and reported sex (48 females, 48 males). Age of the donors ranged between 7 and 75 years old.

#### GTEx Cohort

Complete description of the Genotype-Tissue expression (GTEx) cohort was published previously^72; 73^. In this study, 96 individuals (33 females, 63 males; age range 21-68) with genotype and liver gene expression data were included. 37 of the subjects were organ donors and 59 were postmortem. Liver needle biopsy samples from each subject were obtained in two centers. Samples were preserved in PAXgene tissue kits and shipped to the GTEx Laboratory Data Analysis and Coordinating Center LDACC at the Broad Institute for processing^72; 73^. All GTEx datasets used in this study were from GTEx Analysis ReleaseV6p^27^.

### ChIP-Seq Experimental Protocol

#### Penn Cohort 1

Between 40 and 900 mg of liver wedge biopsies were obtained from each donor prior to transplantation surgery at the Penn Transplant Institute. Flash frozen liver wedge biopsies were processed in a total of 8 batches (six/eight randomized samples per day). On each tissue preparation day, 20 mg of tissue from each liver sample was cut, placed in 1 ml of RNAlater, and flash frozen for isolation of DNA and RNA at a later date. When available 120 mg of tissue from each subject was processed for the Chromatin ImmunoPrecipitation (ChIP) experiment (From 33 subjects, 120 mg of tissue could be used. Tissue amount from the remaining 17 subjects was limited therefore the largest amount available was used; ranging between 20 and 110 mg). The tissue was cut into small pieces (~1 mm^3^), washed with PBS, and fixed with 1% formaldehyde for 5 minutes at room temperature. Nuclei were prepared with the Covaris truChIP™ Tissue Chromatin Shearing Kit with SDS Shearing Buffer according to manufacturer's recommendations. Chromatin was sheared for 14 minutes at 5% duty cycle, 140 Watts peak incident power and 200 cycles per burst using a Covaris S220 Focused-ultrasonicator. Shearing efficiency was assessed using the Agilent High Sensitivity DNA kit and chromatin concentration was determined using a NanoDrop Spectrophotometer. From each subject, a 0.5 ug aliquot of sheared chromatin was kept aside to be used as input chromatin. Each immunoprecipitation was performed using 9 ug of sheared chromatin and 5 ug of antibody (H3K27ac:ab4729, H3K4me3:ab8580) with an overnight incubation at 4°C following the Magna ChIP™ A/G Chromatin Immunoprecipitation Kit protocol. After elution and reverse-crosslinking of Protein-DNA complexes, DNA was cleaned with a QIAGEN QIAquick PCR Purification Kit and quantified using the Agilent High Sensitivity DNA kit. 40 H3K27ac and 45 H3K4me3 samples yielded sufficient DNA (≥2 ng) to generate sequencing libraries. 2 or 5 ng of immunoprecipitated and input DNA was used to generate sequencing libraries using the NEBNext^®^ ChIP-Seq Library Prep Master Mix Set for Illumina. Libraries were sequenced to generate 100 bp single-end reads on Illumina HiSeq2500 instruments at the Penn Next-Generation Sequencing Core.

### RNA-Seq and Genotyping Experimental Protocol

#### Penn Cohort 1

RNA and DNA were extracted in a total of 4 batches (12 or 14 randomized samples per day) using QIAGEN’s AllPrep DNA/RNA/miRNA Universal Kit. Barcoded, strand-specific, polyA+ selected RNA-seq libraries were generated using the Illumina TruSeq Stranded mRNA kit. Quality of each library was assessed using the Agilent Bioanalyzer High Sensitivity DNA Kit. Libraries were then pooled into one group and sequenced to generate 125 bp paired-end reads on Illumina HiSeq2500 instruments at the Penn Next-Generation Sequencing Core. DNA was genotyped using Illumina HumanCoreExome arrays at the Center for Applied Genomics Core at the Children’s Hospital of Pennsylvania.

#### Penn Cohort 2

RNA and DNA extraction method, and library preparation was identical to that of Penn Cohort 1. Libraries were pooled into two groups of 48 samples and sequenced to generate 125 bp paired-end reads on Illumina HiSeq2500 instruments at the Penn Next-Generation Sequencing Core. DNA was genotyped using Illumina HumanCoreExome arrays at the Center for Applied Genomics Core at at the Children’s Hospital of Pennsylvania.

#### GTEx

RNA was extracted from 96 human liver samples as described previously^74^. Non-strand specific, polyA+ selected RNA-seq libraries were generated using the Illumina TruSeq protocol. Libraries were sequenced to generate 76-bp paired end reads. DNA was extracted from whole blood using the QIAGEN Gentra Puregene method and genotyped using the Illumina Human Omni 2.5M and 5M-Quad BeadChip as described previously^74^.

### SPATIaL-Seq Experimental Protocol

#### Cell fixation for chromatin capture

The protocol used for cell fixation was similar to previously published methods^28^. HepG2 cells were collected and single-cell suspension was made with aliquots of 10^7^ cells in 10 ml media (RPMI + 10% FCS). 540 µl 37% formaldehyde was added and incubation was carried out for ten minutes at room temperature in a tumbler. The reaction was quenched by adding 1.5 ml, 1 M cold glycine (4°C). Fixed cells were centrifuged for five minutes at 1,000 g at 4 °C, and supernatant was removed. The pellets were washed in 10 ml cold PBS (4 °C) by centrifugation for five minutes at 1,000 g at 4 °C. Supernatant was removed and cell pellets were resuspended in 5 ml of cold lysis buffer (10 mM Tris pH8, 10 mM NaCl, 0.2% NP-40 (Igepal) supplemented with protease inhibitor cocktails). Resuspended cells were incubated for 20 minutes on ice and centrifuged to remove the lysis buffer. Finally, the pellets were resuspended in 1 ml lysis buffer and transferred to 1.5 ml Eppendorf tubes prior to snap freezing (ethanol/dry ice or liquid nitrogen). Cells were stored at −80 °C until they were thawed again for digestion.

#### 3C library generation

For preparation of initial 3C libraries, 10 million cells were harvested and fixed. The DNA was digested using DpnII, then re-ligated together using T4 DNA ligase and finally isolated by phenol/chloroform extraction. In line with the previously published Capture C protocol^28^, the 3C libraries were utilized for the capture procedure. Cells were thawed on ice, spun down and the lysis buffer was removed. The pellet was resuspended in water and incubated on ice for 10 minutes, followed by centrifugation and removal of supernatant. The pellet was then resuspended with 20% SDS and 1X NEBuffer DpnII and incubated at 37 °C for one hour at 1,000 rpm on a MultiTherm (Sigma-Aldrich). Triton X-100 (at 20% concentration) was added and the pellet was incubated for another one hour. After the incubation, 10 µL 50 U/µL DpnII (NEB) was added and left to digest for eight hours. An additional 10 µL DpnII was added and digestion was left overnight at 37 °C. The next day, another 10 µL of DpnII was added and incubated for an additional three hours. The chromatin was then ligated overnight (8 µL T4 DNA Ligase, HC ThermoFisher (30 U/µL); with final concentration, 10 U/ml) and shaken at 16 °C at 1,000 rpm on the MultiTherm. The next day, an additional 2 µL T4 DNA ligase was spiked in to each sample, and incubated for three more hours. The ligated samples were then de-crosslinked overnight at 65 °C with Proteinase K (Invitrogen) and the following morning incubated for 30 minutes at 37 °C with RNase A (Millipore). Phenol-chloroform extraction was then performed, followed by an ethanol precipitation overnight at −20 °C and then washed with 70% ethanol. Digestion efficiencies of 3C libraries were assessed by gel electrophoresis on a 0.9 % agarose gel and quantitative PCR (SYBR green, Thermo Fisher).

#### SPATIaL-Seq

Custom capture baits were designed using Agilent SureSelect library design targeting both ends of DpnII restriction fragments encompassing promoters (including alternative promoters) of all human coding genes, noncoding RNA, antisense RNA, snRNA, miRNA, snoRNA, and lincRNA transcripts, totaling 36,691 RNA baited fragments. The capture library design successfully covered 95% of the coding gene promoters and 88% of RNA types described above. Custom capture bait design failed for 5% of the coding genes, which were either duplicated genes or contained highly repetitive DNA in their promoter regions. The isolated DNA of the 3C libraries generated by DpnII digestion and ligation were quantified using a Qubit fluorometer (Life technologies), and 10 μg of each library was sheared in dH2O using a QSonica Q800R to an average DNA fragment size of 350 bp. QSonica settings used were 60% amplitude, 30 seconds on, 30 seconds off, 2 minute intervals, for a total of (5 intervals) at 4°C. After shearing, DNA was purified using AMPureXP beads (Agencourt), the concentration was checked via Qubit and DNA size was assessed on a Bioanalyzer 2100 using a 1000 DNA Chip. Agilent SureSelect XT Library Prep Kit (Agilent) was used to repair DNA ends and for adaptor ligation following the standard protocol. Excess adaptors were removed using AMPureXP beads. Size and concentration were checked again before hybridization. 1 ug of ligated library was used and the standard protocol of the SureSelect XT capture kit was followed to obtain the custom designed Capture-C library. The quality of the captured library was assessed using both Qubit fluorometer and Bioanalyzer’s high sensitivity DNA chip. Each SureSelect XT library was initially sequenced on one lane of HiSeq 4000 machine to generate 100-bp paired end reads for QC purposes. All SPATIaL-Seq libraries were then sequenced three at a time on an S2 flow cell on an Illumina NovaSeq machine, generating ~1.6 billion paired-end reads per sample.

### ChIP-Seq Data Processing

#### Penn Cohort 1

Quality of the raw sequence data was assessed using FastQC. Low quality base calls and sequencing adapters were trimmed using Trim Galore! with the following parameters: -stringency 5 -length 50 -q 20. Reads were then aligned to the reference human genome (hg19) using the BWA-MEM algorithm^75^. Aligned reads were sorted and filtered based on a minimum mapping quality of 10 using SAMtools-1.3.1^76^. MACS2^77^ was used to call peaks for each individual ChIP data using the following parameters: --nomodel --extsize 147 -q 0.01 and the corresponding input data as control. Samples that had fraction of reads in peaks (FRiP) ≥1% and that displayed a significant overlap (i.e. right tailed Fisher’s P-value of <10^−6^ and at least two-fold enrichment) with ENCODE DNaseI Hypersensitive sites (the ENCODE DNaseI Hypersensitive site master list generated by the ENCODE Analysis Working Group downloaded in October 2016) were retained for the downstream analyses (Table S1). See Figure S1 for the heatmap plot of Spearman’s correlation of normalized and averaged ChIP-seq read counts for 27 samples that passed the ChIP-Seq QC thresholds. To generate Figure S1, deepTools^78^ was used to normalize the ChIP-Seq read counts to 1x depth of coverage while excluding chromosome X and average scores were calculated based on 10 kb bins that consecutively cover the entire genome.

To define the final set of ChIP-Seq peaks, ChIP-Seq data from biological replicates (N=9 for H3K4me3 and N=18 for H3K27ac) as well as their corresponding input data (N=9 for Input of H3K4me3 and N=18 for Input of H3K27ac) were pooled into separate groups. See Figure S2 for heatmap and profile plots of read density signal around TSS (based on GENCODE v19 transcript annotations) and Figure S3 for correlation between read density signal around TSS and gene expression levels. MACS2^77^ was used to call the peaks on the pooled ChIP-seq data of 9 and 18 individuals respectively while using the corresponding input data as control. Among peaks that were called, 68,600 H3K4me3 and 131,293 H3K27ac peaks that have a mean read count of at least 20 were included in further analyses. See Table S2 for chromosomal positions of the peaks, Figure 2A for distribution of peak lengths and Figure S4 for genomic annotation of ChIP-Seq peaks. Genomic annotations were obtained using Bioconductor’s GenomicFeatures package^79^ and based on GENCODE v19 transcript annotations. H3K4me3 and H3K27ac peaks identified displayed significant enrichment for ENCODE DNase and FAIRE open chromatin regions as well as ENCODE H3K4me3 and H3K27ac sites in HepG2 cells (Figure S5). Links to ENCODE datasets used are included in Table S5. Fisher’s exact test was used to compare the observed numbers of overlaps with ENCODE data sets to the mean overlap of 1,000 sets of randomly selected size-matching regions for each of the original peaks.

### RNA-Seq Data Processing

#### Penn Cohort 1

One outlier sample with fewer than one million reads was excluded from the analysis. Quality of the raw sequence data was assessed using FastQC. Low quality base calls and sequencing adapters were trimmed using Trim Galore! with the following parameters: -stringency 5 -length 50 -q 20 --paired. Trimmed reads were aligned to the reference human genome (hg19) and transcriptome (GENCODE v19 annotations) using STAR v2.5^80^ in 2-pass mode using the following parameters: --outFilterMultimapNmax 10 --outFilterMismatchNmax 10 --outFilterMismatchNoverLmax 0.3 --alignIntronMin 21 --alignIntronMax 0 --alignMatesGapMax 0 --alignSJoverhangMin 5 --twopassMode Basic --twopass1readsN 500000000 --sjdbOverhang 124. Aligned reads were sorted and filtered to retain only primary aligned reads using SAMtools-1.3.1^76^ See Figure S6 for a histogram of number of primary aligned reads. RSEM^81^ was used to estimate gene-level expression as Transcripts Per Million (TPM). 19,133 genes with RSEM expected read count of >6 and TPM of >0.1 in at least 10% of the subjects were defined as expressed. TPM values of the expressed genes were natural log transformed after adding a pseudocount of 1. After log transformation, expression values were quantile normalized between individuals across all expressed genes. For each gene, expression values were then inverse quantile normalized to a standard normal distribution across individuals.

#### Penn Cohort 2

RNA-Seq data of Penn Cohort 2 was processed the same way as in Penn Cohort 1. 19,537 genes with RSEM expected read count of >6 and TPM of >0.1 in at least 10% of the subjects were defined as expressed in this cohort.

#### GTEx

Low quality base calls and sequencing adapters were trimmed using Trim Galore!. Trimmed reads were aligned to the reference human genome (hg19) and transcriptome (GTEx’s gene level model; gencode.v19.genes.v6p_model.patched_contigs.gtf) using STAR v2.5^80^ in 2-pass mode using the same parameters as in Penn Cohorts 1 and 2 except the parameter --sjdbOverhang 75 to correspond to the RNA-seq read length of this cohort. Similarly, aligned reads were sorted and filtered to retain only primary aligned reads using SAMtools-1.3.1^76^. RSEM^81^ was used to estimate gene-level expression as Transcripts Per Million (TPM). 22,415 genes with RSEM expected read count of >6 and TPM of >0.1 in at least 10% of the subjects were defined as expressed.

### Genotype Data Processing

#### Penn Cohort 1

Genotype data was subjected to standard QC checks using whole genome association analysis toolset PLINK^82^. First, genetic sex of individuals was compared to the self-reported sex. One out of 50 subjects had inconsistency between self-reported (male) and genotyped sex (female) but the subject was retained in the study because genotype data was concordant when genotypes based on genotyping array and RNA-Seq data were compared. Next, variants with HWE P-value of <10^−6^ and variants with more than 5% missing rate were excluded. QC’ed genotype data was phased and imputed with SHAPEIT2^83^ and IMPUTE2^84^, respectively, using multi-ethnic panel reference from 1000 Genomes Project Phase 3^85^. Following imputation, variants with HWE P-value of <10^−6^, missing rate of >5%, minor allele frequency (MAF) of <5%, and imputation info score of <0.4 were excluded. This yielded in a total of 4,584,583 imputed variants.

#### Penn Cohort 2

Genotype data of Penn Cohort 2 was processed the same way as in Penn Cohort 1. Two out of 96 subjects had inconsistencies between their self-reported (female) and genotyped sex (male). Both subjects were retained in the study after making sure the genotypes based on genotyping array and RNA-Seq data were concordant. After imputation and QC checks, 4,541,981 variants were retained.

#### GTEx

Genotype data was phased and imputed as described previously^27^. Variants with HWE P-value of <10^−6^, missing rate of <5%, minor allele frequency (MAF) of <5%, and imputation info score of <0.4 were excluded. 5,598,884 variants were retained after imputation and QC filtering.

### SPATIaL-Seq Data Processing

Paired-end reads were pre-processed with the HICUP pipeline^86^, with bowtie2 as aligner and hg19 as reference genome. Significant promoter interactions at 1-DpnII fragment resolution were called using CHiCAGO^29^ with default parameters except for binsize which was set to 2500. To obtain significant interactions at 4-DpnII fragment resolution, artificial. baitmap and ․rmap files were generated where, moving along each chromosome from start to end, consecutive DpnII fragments were grouped by 4 into larger (dijoint) fragments. Significant interactions at 4-DpnII fragment resolution were then called with CHiCAGO using these artificial ․baitmap and ․rmap files and using default parameters except for removeAdjacent which was set to False. Results from the two resolutions were merged by taking the union of the interaction calls at either resolution and removing any 4-fragment interaction which contained a 1-fragment interaction.

### Estimating Population Structure

Principal Component Analysis (PCA) as implemented in EIGENSOFT^87^ was performed using the genotype data of each cohort in aggregate with HapMap Phase 3 genotype data from 1,184 individuals from 11 populations (ASW: African ancestry in Southwest USA; CEU: Utah residents with Northern and Western European ancestry from the CEPH collection; CHB: Han Chinese in Beijing, China; CHD: Chinese in Metropolitan Denver, Colorado; GIH: Gujarati Indians in Houston, Texas; JPT: Japanese in Tokyo, Japan; LWK: Luhya in Webuye, Kenya; MEX: Mexican ancestry in Los Angeles, California; MKK: Maasai in Kinyawa, Kenya; TSI: Toscani in Italia; YRI: Yoruba in Ibadan, Nigeria)^88^. In Penn Cohort 1, 34 of 50 individuals clustered with the HapMap European populations, 12 of them clustered with the HapMap African populations, and the remaining 4 individuals displayed mixed genetic ancestry (Figure S7). In Penn Cohort 2, 62 of 96 individuals clustered with the HapMap European populations, 24 of them clustered with the HapMap African populations, and the remaining 10 individuals displayed mixed genetic ancestry (Figure S8). In GTEx, 81 of 96 individuals clustered with the HapMap European populations, 12 of them clustered with the HapMap African populations, one individual clustered with the HapMap Asian populations and the remaining two individuals displayed mixed genetic ancestry (Figure S9).

### Mapping cis-expression Quantitative Trait Loci (cis-eQTLs)

cis-eQTLs were mapped by linear regression as implemented in FastQTL v2.184^30^. Associations between total expression level (normalized TPM values) of each autosomal gene and variants within 1 Mb of the transcription start site (TSS) were tested within each cohort while adjusting for sex, first three genotype-based PCs, and peer factors. In the GTEx cohort, genotyping platform was additionally included as a covariate in eQTL mapping. The most suitable effective number of Peer factor was determined to be 5, 22, and 16 for Penn Cohort 1, Penn Cohort 2, and GTEx cohorts, respectively (Figure S10). Nominal P-values between each variant and gene pair within 1 Mb of the TSS were calculated by measuring the Pearson product-moment correlation coefficients and using standard significance tests for Pearson correlation^30^. To identify the most significantly associated variant per gene, adjusted P-values were estimated by beta approximation method using the parameter “--permute 10000”. Genome-wide significance was determined by correcting the adjusted P-values for multiple testing across genes using Benjamini&Hochberg method (FDR < 0.05 were considered significant).

METAL^31^ was used for meta-analysis of cis-eQTL mapping by combining nominal P-values across 3 cohorts while taking the sample size and direction of effect into account. For each gene, the most significantly associated variant per gene (i.e. the one with the smallest meta P-value) was recorded to form the empirical, true meta p-value distribution. To assess the significance of meta P-values, eQTL mapping within each cohort was repeated using permuted gene expression data. METAL was successively run on the permuted eQTL results across 3 cohorts and permuted meta P-values were obtained. For each gene, the most significantly associated variant per gene (i.e. the one with the smallest meta P-value) was recorded to form the empirical, null meta P-value distribution. Next, FDR of 0.05 was estimated such that Probability(P-value0<z)/ Probability(P-value1<z)=0.05, where Probability(P-value0<z) is the fraction of P-values from the null meta P-value distribution that fall below the P-value threshold z and Probability(P-value1<z) is the corresponding fraction in the true meta P-value distribution (See Figure S11 for QQ-plots of meta cis-eQTL associations, Figure S12 for relative distance of meta cis-eQTLs to their target gene TSS, and Table S4 for significant meta cis-eQTL results).

### Identification of Shared and Liver-Specific cis-eQTLs

Among 2,625 lead cis-eQTLs, 2,552 of them were tested in GTEx Analysis ReleaseV6p. For each of these 2,552 lead cis-eQTL-gene expression pair, posterior probability of association in 43 non-liver GTEx tissues^27^ were calculated using METASOFT^32^. cis-eQTLs with a posterior probability of 0.9 in at least 38 non-liver tissues were defined as “shared-eQTLs” and cis-eQTLs with a posterior probability of 0.9 in fewer than 5 non-liver tissues were defined as “liver specific eQTLs”. Overlap between these three sets of cis-eQTLs (total, shared, and liver-specific) and H3K4me3 and H3K27ac peaks were identified using bedtools intersect-u function. Liver H3K4me3 and H3K27ac peaks identified in this study as well as those of ENCODE consortium (links to ENCODE datasets are in Table S5) were included in this analysis. For each cis-eQTL set, 1000 matching SNP sets (LD of r^2^ 0.5, MAF of ±5%, Gene density of ±5%, Distance to nearest gene of ±50%, LD buddies of ±50% in European 1000G Phase3) were obtained using SNPsnap^89^. Odds ratios of observed/expected overlaps were plotted in Figure S13 and Figure 3B.

### Mapping trans-expression Quantitative Trait Loci (trans-eQTLs)

Associations between each expressed autosomal protein coding gene and variants that are more than 5 Mb apart were considered as trans. Trans-eQTLs were mapped using MatrixeQTL^90^ while adjusting for the same covariates that were used in cis-eQTL mapping (i.e., sex, ancestry, peer factors in each cohort as well as genotyping platform in GTEx cohort). Trans-eQTL mapping was performed within each cohort using i) all linkage disequilibrium pruned variants (*R*^2^ > 0.5, plink parameters–indep 50 5 2 across cohorts) ii) variants that were identified as cis-eQTLs in this study (2,625 variants based on meta cis-eQTL results) iii) variants that are likely to affect transcription factor activity. Curated list of transcription factors were downloaded from T. Ravasi et al.^91^. Significant meta cis-eQTLs as well as coding variants of this list of curated transcription factors (based on gnomAD release-170228) were included in the analysis.

METAL^31^ was used for meta-analysis of trans-eQTL mapping by combining nominal P-values across 3 cohorts while taking the sample size and direction of effect into account for each type of trans-eQTL approach. Similarly METAL was run on the permuted trans-eQTL results across 3 cohorts and permuted meta P-values were obtained. Permutation based FDR was calculated as explained in the cis-eQTL mapping section above. See Figure S14 for QQ-plots of meta trans-eQTL associations. There were two statistically significant trans-eQTL findings when genome-wide approach was used. However, both of these results were filtered due to presence of genes near trans-eQTL (within 100 kb) with evidence of RNA-seq read cross-mapping due to sequence similarity (Table S12). There were no significant findings when only cis-eQTLs or only variants likely to affect transcription factor activity were tested as trans-eQTLs.

### Mapping cis-histone Quantitative Trait Loci (cis-hQTLs)

Picard Tools’ MarkDuplicates function was used with the ‘REMOVE_DUPLICATES=true’ parameter set to remove duplicate reads from aligned and q10 filtered ChIP-Seq data (initial ChIP-seq data processing and QC steps were explained above in the ChIP-Seq data processing section). Genotype data of Penn cohort 1 was further processed to include only single nucleotide substitutions in hQTL mapping. Allele-specific read counts were obtained using GATK ‘s92 ASEReadCounter function. Total feature counts and GC% values of each feature were used to generate sample specific offset values for each feature. To generate Peer Factors, FPM values (equivalent of TPM for ChIP-Seq data) were calculated, quantile normalized between individuals, and inverse quantile normalized to a standard normal distribution within each peak. Sex, first three genotype-based PCs, five peer factors were included as covariates. RASQUAL^20^ was used to map hQTL associations. Variants within 10 kb of each end of an histone peak were considered as cis. All autosomal peaks were included in hQTL mapping. This corresponded to a total of 65,649 and 128,822 tested peaks for H3K4me3 and H3K27ac, respectively. For each peak, the most significant P-value was selected to form the empirical, true hQTL P-value distribution. To assess genome-wide significance, RASQUAL was successively run using the -r/--random-permutation option. For each peak, the most significant P-value from the permutation run was selected to form the empirical, null hQTL P-value distribution. Next, FDR of 0.05 was estimated such that Probability(P-value0<z)/ Probability(P-value1<z)=0.05, where Probability(P-value0<z) is the fraction of P-values from the null hQTL P-value distribution that fall below the P-value threshold z and Probability(P-value1<z) is the corresponding fraction in the true hQTL P-value distribution. Peaks with significant hQTLs were excluded if they had potential reference mapping bias (ɸ<0.25 and ɸ>0.75). See Table S7 for significant hQTL results.

### Integrative Analyses of Liver cis-hQTLs with Other Functional Datasets

Peaks with cis-hQTLs were overlapped with 61 different transcription factors’ binding sites in HepG2 cells using bedtools intersect –u function (links to ENCODE datasets used are included in Table S5). Enrichment of overlap was calculated relative to 1,000 sets of randomly chosen matching numbers of H3K4me3 and H3K27ac autosomal peaks in our data. One-sided Fisher’s exact test was used to assess the significance of the enrichment.

For 151 of the hQTL-Peaks, target interacting gene promoters could be identified using the SPATIaL-Seq data. For each variant that was located within such hQTL-Peak, cis-eQTL P-value for the interacting gene was pulled to form the distribution of observed cis-eQTL P-values. This observed P-value distribution was then compared to the expected distribution of P-values that was observed when 151 histone peaks with no hQTLs were chosen randomly from the set of autosomal histone peaks that do not have significant hQTLs. In a complementary analysis, the proportion of eQTL-Genes among the 210 genes that interact with hQTL-Peaks was compared with the proportion that was observed among 1,000 sets of randomly chosen 210 genes that were baited, expressed but that do not interact with hQTL-Peaks. Significance was assessed based on the permutation P-value.

### Identification of Shared Genetic Signals Underlying Variation in Histone Modification States and Gene Expression Levels

For each gene with a significant meta cis-eQTL, H3K4me3 and H3K27ac peaks with significant hQTLs that are located within 1 Mb of its transcription start site (TSS) were selected as candidate co-regulated peaks. Gene-Peak pairs with r^2^>0.8 between the lead hQTL and lead eQTL were considered as putatively co-regulated. r^2^ was calculated in the 1000 Genomes, Phase 3, European population. Among significant co-regulation results, the ones in the MHC region (Chr6:28,510,120-33,480,577) were excluded owing to complicated LD patterns of this locus.

### Identification of Causal Genes and Gene Regulatory Regions in GWAS Loci

GWAS summary statistics of twenty phenotypes including coronary artery disease^34^, HDL cholesterol^35^, LDL cholesterol^35^, total cholesterol^35^, triglycerides^35^, diastolic blood pressure^36^, systolic blood pressure^36^, mean arterial pressure^36^, rheumatoid arthritis^37^, Type 2 diabetes^38^, multiple sclerosis^39^, asthma^39^, psoriasis^39^, Parkinson’s disease^39^, Alzheimer’s disease^40^, schizophrenia^41^, Crohn’s disease^42^, ulcerative colitis^42^, inflammatory bowel disease^42^ and age-related macular degeneration^43^ were obtained (See Table S9 for links to data sets). Among GWAS SNPs with P-value < 10^−6^, most significant SNP was chosen per each 1-Mb window. The GWAS locus was then defined as the 2 Mb region around each lead SNP (1Mb upstream and 1 Mb downstream). Approximate Bayes Factor colocalization analysis of the coloc package^44^ was performed between the GWAS trait and each gene with significant meta cis-eQTL whose TSS was in the GWAS locus. Among colocalization pairs that pass the colocalization power threshold (PP3+PP4>0.8), the ones that met PP4/(PP3+PP4)>0.9 threshold were considered as significantly colocalized. Similarly, significant hQTL-Peaks that were located within each GWAS locus were selected as candidate co-regulation pairs. GWAS-Peak pairs with r^2^>0.8 between the lead hQTL and lead GWAS SNP were considered as putatively co-regulated. r^2^ was calculated in the 1000 Genomes, Phase 3, European population. All significant co-regulation and colocalization signals in the MHC region (Chr6:28,510,120-33,480,577) were excluded owing to complicated LD patterns of this locus.

### Identification of Candidate Causal Variants

For each candidate causal gene and gene regulatory region identified in GWAS loci, approximate Bayes factors (ABF) were estimated using the approach previously described^60; 61^. Briefly, the original eQTL/hQTL P-value of each SNP was converted to a standard one-tailed Z-score using inverse normal cumulative distribution function. To compute the ABF, the standard deviation for the prior was set to 0.15 and the variance of the effect estimate, *V*, was approximated using the MAF and sample size. For each candidate gene and gene regulatory region, the smallest set of variants that account for 95% of the posterior probability was calculated. Loci with fewer than 5 candidate causal variants are shown in Figure 7A.

## DESCRIPTION OF SUPPLEMENTAL DATA

Supplemental Figures include Figures S1-S14. Supplemental Tables include Tables S1-S12.

## ACKNOWLEDGEMENTS

We thank all liver donors and their families for participating in this study. We thank Ana Gonzalez and Mary Shaw of Penn Transplant Institute for coordinating donor tissue and donor information sharing. We thank Daniel Savic for his input on ChIP-Seq experiments, Jonathan Schug, and Susannah Elwyn for their help with generating RNA-Seq and ChIP-Seq data, Arif Harmanci for his input in ChIP-Seq peak calling, Graham McVicker for his input on generating read density plots, and Ben Strober for sharing trans-eQTL cross-mapping regions list. We thank Greg Cooper, Barbara Engelhardt, and Katalin Susztak for helpful discussions and comments on an earlier draft of the manuscript. This study was supported by NIH grants R01 HL133218 (C.D.B.), R01 AG057516 (S.F.A.G.), HL055323 (A.M.B., D.J.R.), HL134853 (N.J.H., D.J.R.), HL109489 (D.J.R.) and HG006398 (D.J.R.). The Genotype-Tissue Expression (GTEx) Project was supported by the Common Fund of the Office of the Director of the National Institutes of Health, and by NCI, NHGRI, NHLBI, NIDA, NIMH, and NINDS. M.Ç. was partially supported by Penn IBI Postdoctoral Collaboration Fellowship. S.F.A.G is supported by the Daniel B. Burke Endowed Chair for Diabetes Research.

## ACCESSION NUMBERS

Genotype, RNA-Seq, and ChIP-Seq datasets from Penn Cohorts have been deposited in the dbGaP under ID code of <tbd>. Genotype and RNA-Seq datasets from GTEx Cohort have been deposited in the dbGaP under ID code of phs000424.v6.p1. HepG2 SPATIaL-Seq data have been deposited in ArrayExpress (www.ebi.ac.uk/arrayexpress/) with accession number E-MTAB-7144.

